# Functional brain imaging predicts population-level visits to urban spaces

**DOI:** 10.1101/2024.02.16.580650

**Authors:** Ardaman Kaur, André Leite Rodrigues, Sarah Hoogstraten, Diego Andrés Blanco-Mora, Bruno Miranda, Paulo Morgado, Dar Meshi

## Abstract

Urbanization is increasing around the world, and urban development strategies focusing on sustainability and the welfare of urban residents are needed. In response to this need, the field of neurourbanism has emerged, which leverages research on the human brain to understand and predict the influence of urban environments. For example, studying brain regions involved in reward processing and value-based decision making, such as the ventromedial prefrontal cortex (vmPFC), may help us understand how people interact with and navigate through urban environments. In this study, we aimed to ascertain whether neural activity within the vmPFC can predict population-level visits around the urban spaces of a city – in our case, Lisbon, Portugal. We used the density of photographs taken around Lisbon as a proxy measure of these visits. To do this, we created a stimulus set featuring 160 images of Lisbon sourced from the social media platform, Flickr. Then, study participants in the U.S. who had never visited Lisbon, viewed these images while we recorded their brain activity. We found that in our sample, activity in the vmPFC predicted the density of photographs taken around Lisbon, and hence, the population-level visits. Our research highlights the crucial role of the brain, especially reward-related brain regions, in shaping human behavior within urban environments. By shedding light on the neural mechanisms underlying urban behavior in humans, our research opens exciting possibilities for the future of urban planning. With this knowledge, policymakers and urban planners can potentially design cities that can promote well-being, social interaction, and sustainable living.

## Introduction

Urbanization, the process by which cities and towns grow in dimension and number of inhabitants, is rapidly advancing globally and currently stands at 57%^1^. This shift in urban-area demographics has important implications for overall well-being ^2–5^, as both positive and negative effects have been observed. On the positive side, urban areas generally offer better access to educational and employment opportunities, advanced healthcare facilities, and diverse cultural and recreational activities, which help to contribute to a higher standard of living^6^. However, the swift pace of urbanization also gives rise to formidable challenges attributed to several factors, such as lack of green spaces, increased traffic noise, and varied social inequalities^7–9^. This emphasizes the need to adopt sustainable strategies for urban development, placing a strong emphasis on prioritizing the health, safety, and well-being of all residents.

In this context, the nascent field of neurourbanism has emerged, which leverages measuring the human brain to understand and predict the influence of urban environments on behavior^10^. Neurourbanism has the potential to contribute significantly to the design of cities that prioritize cognitive, emotional, and physical well-being, thereby fostering a shift towards a more human-centric urban design^11–13^. By prioritizing the well-being of individuals, cities can create environments that are conducive to the overall health and happiness of their inhabitants. Neurourbanism capitalizes on brain-scanning tools, such as functional magnetic resonance imaging (fMRI). To explain, fMRI detects changes in blood oxygen levels to identify the brain regions that are active during specific tasks or mental processes. When neurons become active, they require more oxygen, so blood flows to the active areas to meet this demand – fMRI detects the locations where these changes in blood oxygenation levels occur^14^. Conducting fMRI research is costly, but offers the unique ability to image deep brain regions with high spatial resolution. Therefore, the integration of fMRI methodology in neurourbanism research holds promise for informing evidence-based urban planning and design^15^.

Recent advances in fMRI research suggest that specific brain regions can act as neural predictors of behavior, actually having the potential to forecast collective behavior at the population level^16,17^. For example, Falk and colleagues evaluated activity in a brain region called the ventromedial prefrontal cortex (vmPFC) among 33 heavy smokers who had expressed an intention to quit^18^. The participants had their brain activity measured with fMRI while they watched three different anti-smoking television campaigns promoting the National Cancer Institute’s telephone “Quitline” to help smokers quit (1-800-QUIT-NOW). This study’s importance lies in exploring how activity in the vmPFC – a key region in the brain’s reward system involved in valuation and decision-making^19–21^ – could predict the efficacy of these anti-smoking campaigns. Importantly, these same advertisements that the participants watched were also previously broadcast on television, and the researchers compared the Quitline call volume from the month before and the month after each advertisement aired. The study found that the anti-smoking campaign generating the highest Quitline call volume elicited the highest level of neural activity in the vmPFC. This suggests that the neural activity observed in the vmPFC brain region of 33 participants accurately predicted the effectiveness of anti-smoking ads at the population level. In addition to this smoking aspect, the vmPFC is generally involved in valuation and decision making, so activity in this region doesn’t just predict anti-smoking campaign effectiveness. For example, previous research has used reward system activity to predict population-level behavior concerning market reactions to music clips, microloan appeals (requests for small-scale financial assistance), crowdfunding proposals, stock market prices, and food choices^22–28^. Therefore, the brain’s reward system, including the vmPFC, can predict human behaviors across a range of situations, potentially making it a valuable brain region to study with respect to urban planning.

With the above in mind, we considered people’s visits to urban spaces, such as people’s preference for certain areas. We hypothesized that we could measure a small group of participants’ vmPFC response to images of an urban area, and that this brain activity could predict the real-world, population-level visits around this urban area. To address our hypothesis, we created a stimulus set comprising 160 geotagged photos of urban spaces of the city of Lisbon, Portugal from the online photo management and sharing application Flickr^29^. Importantly, we were able to ascertain the density of these photographs taken in each region of the city (see Methods). We then presented this stimulus set to study participants in the U.S. who had not previously visited Lisbon and measured their brain activity using fMRI. We then analyzed our data, determining if neural activity in the vmPFC predicted the density of these photographs taken in locations around the city. If our hypothesis is correct, this study would serve as a foundation and proof of concept for integrating a neuroscientific approach into urban planning, marking a significant step forward in the field of neurourbanism. As we delve further into this interdisciplinary realm, the insights gained from studies such as this can play a pivotal role in transforming urban landscapes to foster health and well-being.

## Methods

### Participants

We used an online platform^30^ to recruit 80 healthy, right-handed students from a large university in the Midwest United States. All participants either earned three course credits or received $50 compensation for taking part in the study. Three participants were unable to complete the experiment due to discomfort in the MRI scanner and were subsequently excluded from the analysis, resulting in a final sample size of 77 (41 males, mean age=20.8 years, SD=2.2). All participants had no history of psychiatric or neurological disorders. No participant had previous in-person exposure to the images viewed in the MRI scanner (participants had never been to Lisbon, Portugal; see below). Our study was approved by the university’s Institutional Review Board, and all participants provided written informed consent.

### Procedure

On the day of the experiment, participants reviewed a detailed document that explained the experimental procedure. They then provided informed written consent and were placed into the MRI scanner. During the imaging session, participants completed our urban environment fMRI task. In this task, participants were presented with images of the urban environment as stimuli and were asked to rate how much they liked the place depicted in each image. Detailed information regarding the stimulus set used in the fMRI task and the task paradigm are explained below.

### Stimulus Set

We created a rich stimulus set of 160 geotagged urban space images of Lisbon, Portugal, sourced from the Flickr social media platform. The complete procedure for creating our stimulus set images is described in a previously published manuscript^31^. Briefly, we extracted bulk geotagged urban photographs (N=75,233) from the Flickr platform taken in the city of Lisbon from January 1, 2016, to September 29, 2021. We divided the entire search area of Lisbon into smaller cells measuring 100 x 100 meters. With the geotag data from each image, we extracted the total number of images in each 100 x 100 meter cell. This allowed us to calculate a cell image density for each cell, which indicates the total number of geotagged photographs shared by Flickr users in that area during the period of interest. We then linked this cell image density with each image taken within that cell (e.g., if image 1 came from a cell with 500 pictures taken within it during the time period, then the cell image density linked to image 1 was 500). To ensure that our stimulus set was representative of urban spaces in Lisbon, we meticulously selected (with an algorithm and visual inspection) 160 images from the larger pool of photographs. Further details about the selection process can be found in the manuscript. The cell image density values of 160 images thus served as a proxy for measuring people’s visitation patterns around the city. To note, past research has established geotagged social media data as a valuable source for gaining insights into a place’s popularity and visitation patterns over time^32–36^. Additionally, we ascertained all brightness and contrast values for each image. To note, after generating the image set and collecting neuroimaging data, we identified three image outliers with values all above three SD from the image set mean: two in cell image density and one in brightness. As a result, even though all 160 images were presented to the participants in the scanner, we excluded the three images associated with the outliers from the fMRI analysis.

### fMRI Task Paradigm

The task paradigm (see Figure 1) consisted of a jittered fixation cross (mean=1.8 s; range=1 to 5s) at the beginning of each trial. Next, participants saw an image of an urban space from the stimuli set and rated it on a scale of one to four based on how much they liked the place portrayed in the image. Participants were able to record their preference ratings using an MRI-compatible response pad, which involved pressing a button ranging from “1=Not at all” (thumb) to “4=Very much” (ring finger). After rating for preference, participants’ responses were immediately displayed by greying out the according number on the display. Participants viewed the image for four seconds total in each trial. The order in which the images were presented was randomized across participants.

**Figure 1.**
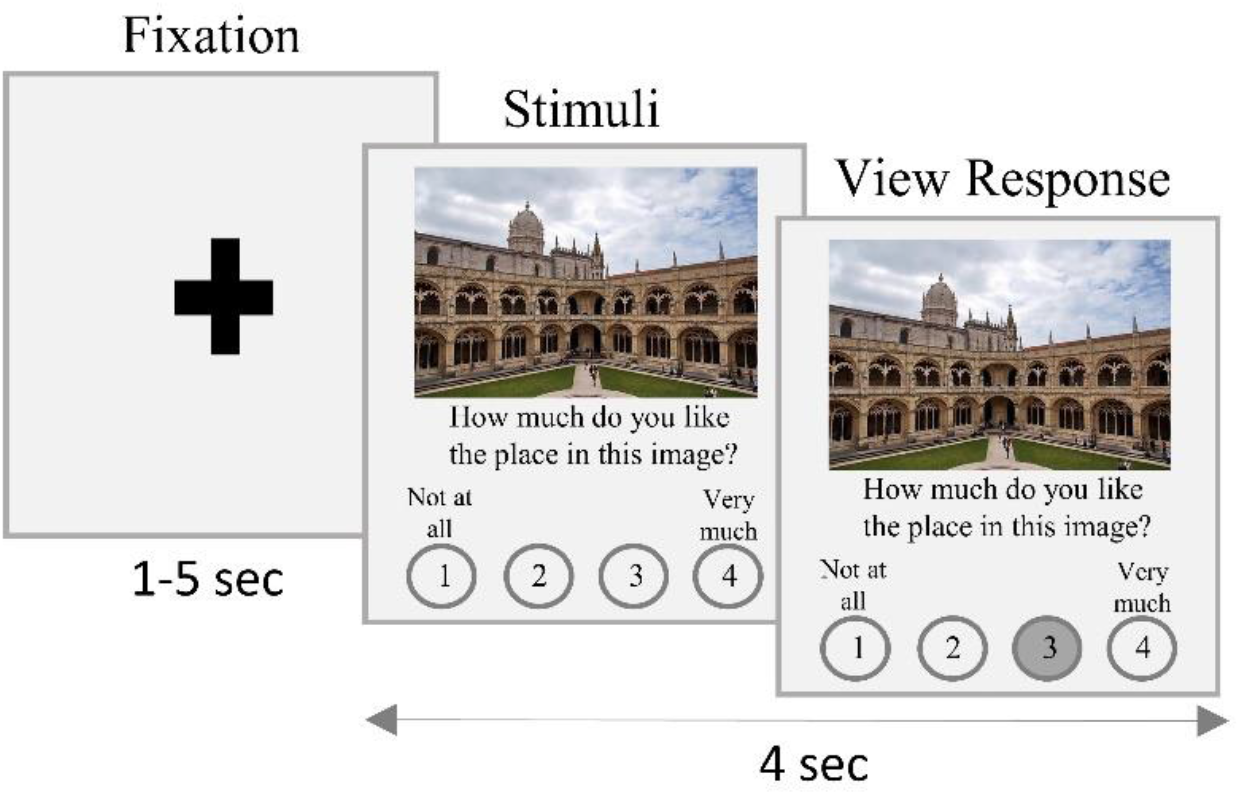
Urban environment fMRI task. At the start of each trial, participants saw a fixation cross, displayed for various durations across trials (mean= 1.8 s; range=1 to 5s). Next, participants saw an image of an urban space in Lisbon, Portugal and rated it on a scale of one to four based on how much they liked the place portrayed in the image. After rating, participants’ responses were immediately displayed by greying out the according number on the display. Participants viewed the image for four seconds total in each trial. Overall, the task consisted of 160 trials, each with a unique image of an urban space.

### MRI Data Acquisition

Whole-brain imaging was performed using a 3T GE scanner (GE HealthCare, United States) with a 16-channel head coil. Anatomical images were acquired using a T1-weighted mprage protocol (256 × 256 matrix, 184 sagittal slices of 1 mm thickness). Blood-oxygen-level-dependent (BOLD) sensitive T2*-weighted functional images were acquired using a single shot gradient-echo imaging pulse sequence (repetition time=2200 ms, echo time=25 ms, flip angle=78°, field of view =220 mm, distance factor =0%, 38 axial slices, 3 × 3 × 3 mm^3^ voxels, interleaved slice ordering). Participants completed the task in two fMRI runs, each with 218 volumes lasting approximately 8 minutes. Stimuli were presented using the Psychtoolbox-3^37^ for MATLAB 2023a (The MathWorks Inc.) on an MRI-compatible screen.

### MRI Data Analysis

We first preprocessed the fMRI data using the preprocessing pipeline for volume-based analysis in CONN software^38^. This preprocessing procedure included the removal of the first four scans (dummy scans) involving significant signal changes. Next, we performed the realignment and unwarping of functional images where all scans were coregistered and resampled to the first scan of the first session using b-spline interpolation. This step also corrects susceptibility distortion by motion interactions. Following the realignment and unwarping step, we translated the functional center to the (0, 0, 0) coordinate and performed the slice time correction of functional data. Subsequently, functional outlier detection (ART-based identification of outlier scans for scrubbing) was performed, where potential outlier scans were identified from the observed global BOLD signal and the amount of subject motion in the scanner. Acquisitions with global BOLD signal changes above five standard deviations or framewise displacement above 0.9 mm were flagged as potential outliers. Functional and anatomical scans were then normalized into standard MNI space and segmented into grey matter, white matter, and CSF tissue classes using segmentation and normalization procedures. Lastly, we carried out functional data smoothening with a Gaussian kernel with an FWHM of 6 mm.

To test our hypothesis regarding the link between brain activity and cell image density, we conducted statistical analyses on individual subject spaces using a General Linear Model (GLM) in SPM12. fMRI time series data were thus modeled as a series of impulses convolved with a canonical Hemodynamic Response Function (HRF). The following regressors were included in our GLM:

1. Fixation regressor: time periods when participants viewed the fixation on the screen.
2. Stimulus viewing regressor: time periods when participants viewed the images of urban spaces from Lisbon on the screen.
3. Stimulus viewing X Parametric modulator 1 regressor: cell image density values for the images.
4. Stimulus viewing X Parametric modulator 2 regressor: preference rating values for the images.
5. Response regressor: time periods after participants pressed a button to rate an image until the end of that trial (for instance, if a participant pressed the button three seconds after the image appeared, the regressor models the duration between the button press and the completion of four seconds, which would be one second).
6. Error regressor: time periods of complete trials when participants didn’t respond or provide preference ratings after the end of the trial and for the images with outlier values for cell image density and brightness/contrast.

The motion parameters for translation (i.e., x, y, and z) and rotation (i.e., yaw, pitch, and roll) were included as regressors of no interest in the GLM. Importantly, to mitigate the potential confounding influence of participant preference ratings, we incorporated their preference ratings as regressors in our GLM (see regressor 4 above), orthogonalizing it to the other parametrically modulated regressor weighted by cell image density (see regressor 3 above). Furthermore, we conducted a separate correlation analysis with the MATLAB-based robust correlation toolbox^39^ to ascertain whether any significant correlations existed between cell image density and the average preference ratings for each image. The observed correlations were weak and statistically non-significant (see Results) ensuring that our analysis accurately disentangles the influence of participant preference from the cell image density measure.

Next, contrast images at the individual level for contrasts, *Parametric modulator 1 (cell image density) > Fixation,* and *Parametric modulator 2 (preference ratings) > Fixation* were computed and taken to a group-level analysis using voxel-wise one-sample t-tests. We performed the group-level analysis within a functional mask for the positive effects of subjective value (SV) on BOLD^40^ (including vmPFC). This functional mask was derived from a meta-analysis of 206 studies of SV that identified key regions of the brain’s reward system in adults. Our selection of this functional mask was guided by our hypothesis, which emphasizes the crucial involvement of the brain regions representing SV in the context of human decision-making and predicting people’s visits around a city and the resulting density of photographs captured in this urban space.

We performed the group-level analysis in CANLAB’s robust regression toolbox^41^. The robust regression toolbox relies on an iteratively reweighted least squares (IRLS) approach to detect and manage influential extreme outliers^42^. This methodology serves the purpose of decreasing the risk of false-positive and false-negative findings while preserving statistical power. Notably, IRLS is beneficial in scenarios with small sample sizes (e.g., n=10), and its advantages become more pronounced as the sample size grows (e.g., n=40) – given our sample size of 77, it can be considered even more advantageous. Furthermore, IRLS effectively controls false-positive rates, even when genuine effects may be absent. This ensures the reliability and validity of statistical inferences. For the masked group-level analysis, the contrast images for contrasts, *Parametric modulator 1 (cell image density) > Fixation, and Parametric modulator 2 (preference ratings) > Fixation* were subjected to voxel-wise False Discovery Rate (FDR) correction, with a threshold of p<0.05.

## Results

### Behavioral Correlations

We examined the relationships between cell image density and average preference ratings by our participants for each image using three different correlation measures: Pearson’s correlation coefficient (r=0.149, p >0.05), Percentage bend correlation (r=0.145, p > 0.05), and Spearman’s correlation (r=0.140, p >0.05). All correlations observed were weak and statistically insignificant, revealing no direct relationship between the average preference ratings provided by individuals inside the scanner and the density of photographs taken in the urban environment. The lack of relationship between these variables is important for the interpretation of the below neuroimaging analysis.

### Neuroimaging Analysis

We examined how the neural activity in reward-associated brain regions, including vmPFC, related to the cell image density of our stimulus set with the *Parametric modulator 1 (cell image density) > Fixation* contrast. Our analysis revealed a significant activation cluster in bilateral vmPFC (left/right MNI coordinates (vx): 104, 211, 84; t = 2.2, p < 0.05, FDR-corrected; see Figure 2 & Table 1). In other words, the vmPFC of a small sample of individuals who have never been to a city can predict the likelihood of a completely different, population-scale group of people going to an urban area and taking/sharing photographs in that urban area. Other brain regions that were significantly associated with cell image density were the bilateral posterior cingulate cortex, bilateral thalamus, left amygdala, and left hippocampus (see Table 1).

**Figure 2.**
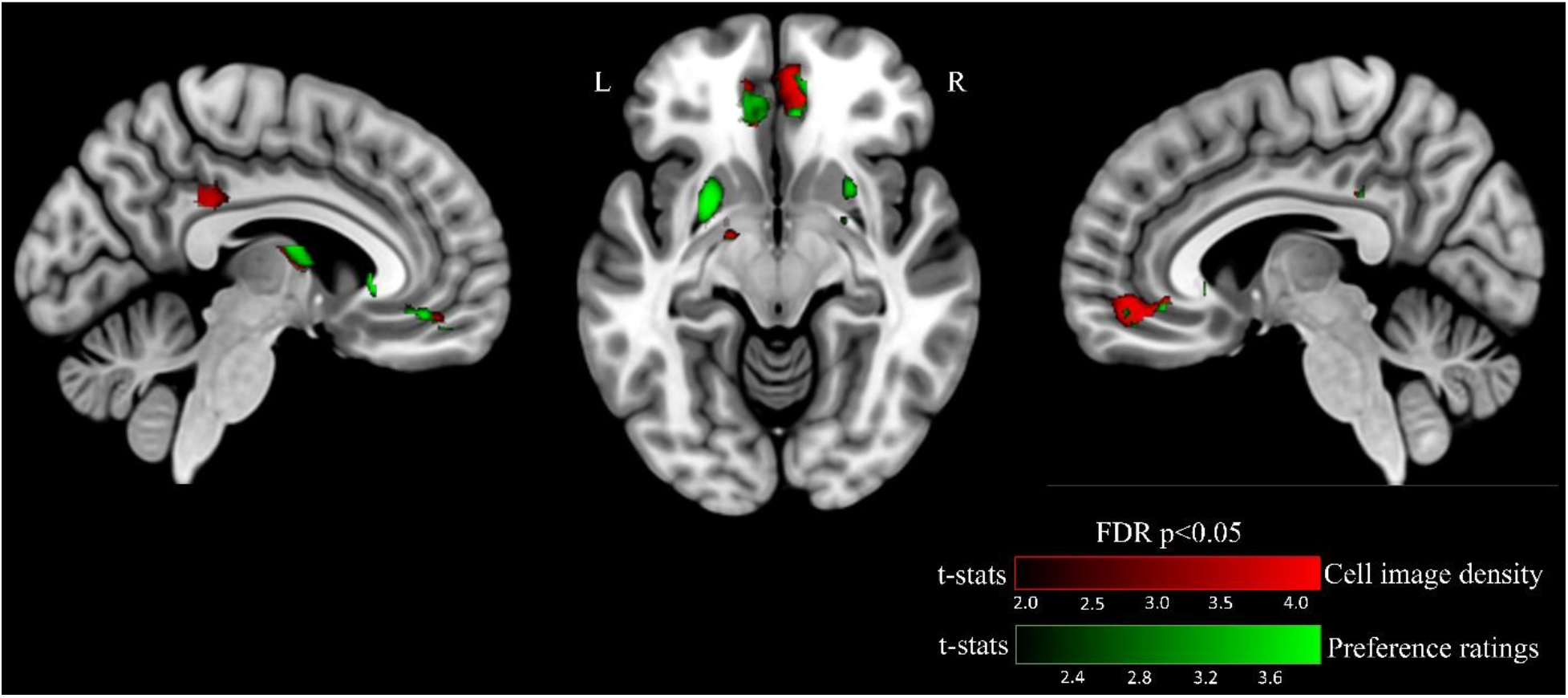
Brain regions related to cell image density and participant preference ratings. Brain activation associated with the trial-by-trial cell image density (number of images in each 100×100 meter cell division of the city of Lisbon, Portugal) in red, and trial-by-trial participant preference ratings provided within the MRI scanner in green. The axial and sagittal slices demonstrate distinct and overlapping significant activations in the ventromedial prefrontal cortex (vmPFC) for cell image density and participant preference ratings at p<0.05 (FDR corrected). Color bars represent t-statistic values. L, left; R, right. Activation is shown in *a priori* defined regions (see Methods).

**Table 1.**
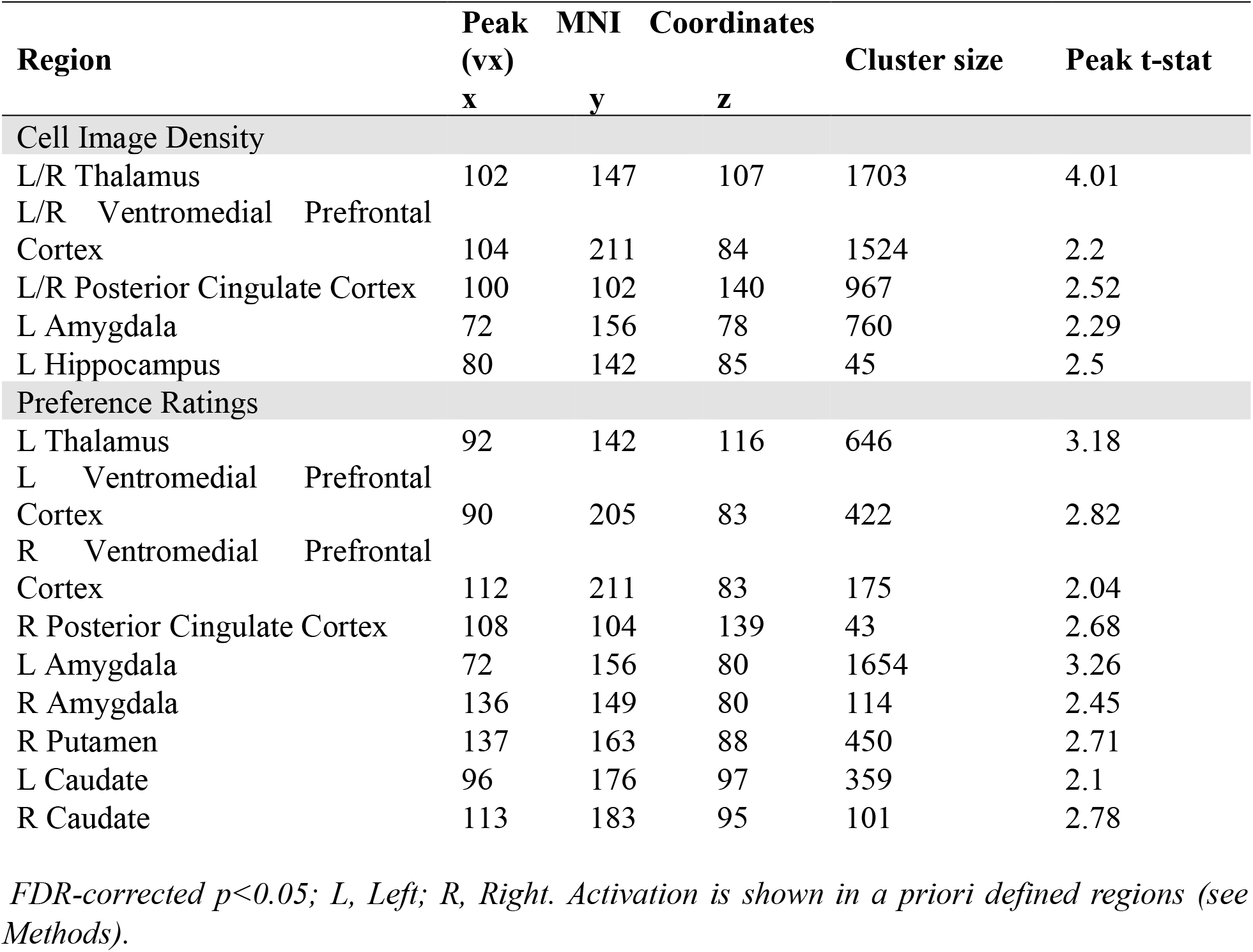
Significant clusters associated with cell image density and preference ratings.

As mentioned in the Methods section, we wanted to tease apart brain activity due to participant image preference from brain activity due to cell image density. Therefore, we added participant preference ratings as a regressor in the same GLM. The *Parametric modulator 2 (preference ratings) > Fixation* contrast from the same model also yielded a significant activation cluster in bilateral vmPFC (left MNI coordinates (vx): 90, 205, 83; t = 2.82, p < 0.05, FDR-corrected; right MNI coordinates (vx): 112, 211, 83; t = 2.04, p < 0.05, FDR-corrected) along with activations in bilateral amygdala and caudate, right posterior cingulate cortex, right putamen, and left thalamus (see Figure 2 & Table 1). Figure 2 illustrates the partially overlapping results for both the *Parametric modulator 1 (cell image density) > Fixation* and *Parametric modulator 2 (preference ratings) > Fixation* contrasts.

## Discussion

This study aimed to determine whether the neural activity in the vmPFC of a small group of individuals could predict population-level visits around a city. We employed a proxy for these population-level visits, which was the density of photographs taken around the city. Seventy-seven participants who had never visited Lisbon underwent fMRI scanning while viewing and rating photographs depicting urban environments in Lisbon, Portugal. We found a significant correlation between the vmPFC activity in the group of fMRI-scanned individuals and the density of photographs (cell image density) taken around Lisbon. Given the vmPFC’s established role in processing value^40^, these findings suggest that neural activity in the vmPFC reflects the valuation of areas in an urban environment, predicting people’s visitation patterns to or away from areas in that environment. In other words, people typically make decisions to maximize value, and because certain urban areas induce greater value-related brain activity than others, people in the city are more likely to travel to, and take pictures of, those urban areas. Notably, our results align with prior studies demonstrating that vmPFC activity can accurately predict real-world aggregated behaviors^11,16,18,23,27,43^, emphasizing this brain region’s crucial role in value computations and the weighing of potential outcomes in decision-making. To the best of our knowledge, the current study is the first to utilize brain activity for the forecasting of population-level movement around physical space, offering potential applications in urban planning and urban design, supporting the overall goals of neurourbanism.

In this study, vmPFC activity not only correlated with the density of photographs but also with individuals’ preference ratings for each urban environment photograph. However, no significant correlation was found between participant preference ratings and cell image density for urban spaces alone, demonstrating that these variables are independent of one another. Furthermore, when participant preference ratings and cell image density were jointly analyzed in our GLM (each with an orthogonalized regressor), selective voxels in vmPFC exhibited a correlation with each regressor. We speculate that our findings suggest the vmPFC to be selective not only for subjective, aesthetic valuation (e.g., participants’ preference ratings), but also for other valuation aspects that could be driving individuals to visit urban spaces. For example, people visit urban spaces because they have socially or culturally relevant value, not just aesthetic value (e.g., a historically relevant building that is not aesthetically pleasing). This implies that individuals may choose to visit and capture photographs of specific places not solely based on personal preferences or interests, but they are likely driven by some common (either perceptual, cognitive, social, or cultural) value associated with those places. Therefore, the predictive capacity of the vmPFC for the density of photographs in urban spaces underscores its role in facilitating behavior change within a sociocultural context, not just an aesthetic one. This aligns with previous research suggesting that the vmPFC is involved in integrating normative influence and individual values and beliefs during behavioral influence^44,45^. Our study highlights the interplay between neural processes in vmPFC, social dynamics, and cultural significance in shaping human behavior in urban environments. Future research can help tease apart contributions from aesthetic values and social or cultural values in motivating people to move around a city.

Alongside the vmPFC, our study also revealed that activity in brain other regions, including the posterior cingulate gyrus, amygdala, and thalamus, also predicted the population-level density of photographs taken in urban areas. These brain regions are not well established to demonstrate the forecasting ability of regions such as the vmPFC, or other well-known reward regions such as the nucleus accumbens. However, research has demonstrated that these brain regions can be involved in value-based decision-making processes. For example, the posterior cingulate gyrus, functioning as a central hub of the brain with extensive connections, has been found to play a role in tracking decision salience, by focusing on deviations from standard choices^46,47^. Moreover, the amygdala and thalamus have also emerged as influential in decision-making, emotions and social behavior, by signaling reward, punishment, and social cues^48,49^. For example, a study by Genevsky and colleagues demonstrated that models incorporating average brain activation in the vmPFC, ventral striatum, amygdala, insula, and inferior frontal gyrus successfully predicted crowdfunding project outcomes^24^. Thus, we believe that future research can further elucidate the specific roles of these brain regions in decision-making within urban settings and, notably, in predicting outcomes like patterns of visitation around a city. A clearer understanding of the neural mechanisms in such complex real-world scenarios is imperative for advancing our comprehension of urban planning and decision-making processes, such as navigation around a city.

It is important to acknowledge a limitation in our current study. Our urban image stimulus set is static in nature, with images selected from a specific time frame (2016 to 2021). Therefore, the static nature of this stimulus set may not fully capture the dynamic and evolving complexity of urban settings. Future research could explore avenues to incorporate dynamic data sources to address these limitations and advance the field. For example, by embracing community-based geoportals and citizen science spatial data infrastructures, researchers can access real-time and continuously updated information, thereby capturing the ongoing evolution of urban landscapes. Integrating dynamic data sources into future research will enhance the adaptability of forecasting models, allowing them to better respond to unforeseen events and changing urban contexts. This integrated framework holds the potential to serve as a valuable tool for both comprehending and forecasting human engagement in various environments, ensuring that urban planning strategies remain adaptable and responsive to the ever-changing nature of urban life.

## Conclusion

This study assessed whether activity in the vmPFC of a small group of individuals could predict population-level visits around a city. To do this, we curated a stimulus set of 160 photos of urban spaces in Lisbon sourced from the social media platform, Flickr. The study’s participants (who had never visited Lisbon) viewed these photographs in an MRI scanner and rated them based on preference. We then conducted an fMRI analysis to explore the relationship between vmPFC activity and the density of the photographs (cell image density) associated with each photo in the stimulus set. Our results revealed that activity in the vmPFC could predict the density of photographs in urban areas, and hence people’s visits to these urban areas. We also established that the neural activity within the vmPFC for the density of photographs was distinct in contrast to the vmPFC voxels that correlated with participants’ average preference ratings for each photograph. In sum, our findings underscore the selectivity of the vmPFC, not only for subjective valuation but also for broader considerations, including social and cultural importance.

Our study’s results contribute to the field of neurourbanism, as well as urban planning and urban design, by shedding light on how our brains may encode information related to the urban environment and drive visits around this environment. Such insights can potentially aid in the development of future human-centric cities, specifically tailored to optimize how our brains perceive and interact with the environment. By incorporating these findings into the urban infrastructure and services, we may be able to create cities that are more efficient and enhance population livability and movement in those urban areas. This neurourbanistic approach can offer valuable insights into specific urban features, informing the development of environments that align with different neural responses. Overall, this approach can contribute to refining urban planning strategies, ultimately fostering enhanced urban livability and well-being.

